# Systematic identification of a CDYL1-dependent decrease in lysine crotonylation at DNA double-strand break sites functionally uncouples transcriptional silencing and repair

**DOI:** 10.1101/2021.05.31.446417

**Authors:** Enas R. Abu-Zhayia, Feras E. Machour, Laila A. Bishara, Bella M. Ben-Oz, Nabieh Ayoub

**Author notes:** Corresponding author: Prof. Nabieh Ayoub, Faculty of Biology, Technion - Israel Institute of Technology, Haifa 3200003, Israel. These authors contributed equally to this work (Feras E. Machour, Laila A. Bishara).

## Abstract

Previously, we showed that CDYL1 is recruited to DNA double-strand breaks (DSBs) to promote homology-directed repair (HDR) and foster transcriptional silencing. Yet, how CDYL1 elicits DSB-induced silencing is not fully understood. Here, we systematically identify a CDYL1-dependent local decrease in the transcriptionally active marks lysine crotonylation (PanKcr) and crotonylated histone residue H3K9cr at *Asi*SI-induced DSBs, which correlates with transcriptional silencing. Mechanistically, we reveal that CDYL1 crotonyl-CoA hydratase activity counteracts PanKcr and H3K9cr at *Asi*SI sites, which triggers the eviction of the transcriptional elongation factor ENL and foster transcriptional silencing. Furthermore, genetic inhibition of CDYL1 hydratase activity blocks the reduction in H3K9cr and alleviates DSB-induced silencing, while HDR efficiency unexpectedly remains intact. Therefore, our results functionally uncouple the repair and silencing activity of CDYL1 at DSBs. In a broader context, we address a long-standing question concerning the functional relationship between HDR and DSB-induced transcriptional silencing, suggesting that they may occur independently.

**Highlights:**

- Systematic identification of a local decrease in lysine crotonylation PanKcr and H3K9cr at *Asi*SI-induced DSBs that correlates with transcriptional silencing.
- CDYL1 crotonyl-CoA hydratase activity downregulates Kcr at DSBs.
- Kcr reduction at DSBs promotes ENL eviction and fosters transcriptional silencing.
- CDYL1 roles in DSB-induced transcriptional silencing and HDR are functionally uncoupled.

## Introduction

Eukaryotic cells encounter thousands of endogenous and exogenous DNA lesions per day that jeopardize genomic stability and may incite carcinogenesis. Among these lesions, double-strand breaks (DSBs) are one of the most detrimental DNA lesions and their improper repair has deleterious consequences including apoptosis, senescence, or oncogenic transformation. To repair these lesions and preserve genome integrity, cells have evolved highly coordinated DNA damage response (DDR) networks (Blackford and Jackson, 2017; Caron and Polo, 2020; Jeggo et al., 2016; Ochs et al., 2019; Tubbs and Nussenzweig, 2017). Eukaryotic cells employ four known DSB repair pathways including homology-directed repair (HDR), classical nonhomologous end joining (c-NHEJ), alternative NHEJ (alt-NHEJ), and single-strand annealing (SSA). While c-NHEJ, alt-NHEJ and SSA are error-prone and can occur throughout the cell cycle, HDR is restricted to S and G2 phases when a sister chromatid is available and serves as a template for scarless and accurate repair of DSBs. The choice between the DSB repair pathways is determined by the chromatin state at DSB sites, DSB-end resection and cell cycle phase (Aleksandrov et al., 2020; Ceccaldi et al., 2016; Clouaire and Legube, 2015, 2019; Clouaire et al., 2018; Ferrand et al., 2021; Hustedt and Durocher, 2016; Jasin and Rothstein, 2013; Scully et al., 2019; Shrivastav et al., 2008).

DSBs are grouped into two main categories: physiological DSBs (also referred as programmed DSBs) and exogenous DSBs (also referred as unprogrammed DSBs). While physiological DSBs activate gene expression, exogenous DSBs induce local and transient transcriptional silencing (Capozzo et al., 2017; Caron et al., 2019; Khan and Ali, 2017; Machour and Ayoub, 2020; Purman et al., 2019; Silva and Ideker, 2019; Ui et al., 2020). ATM, DNA-PK and PARP1 are key determinants of DSB-induced silencing that choreograph the activity of various silencing factors that either directly affect RNA polymerase II (Pol II) activity or modulate chromatin structure at DSB sites (Machour and Ayoub, 2020). For instance, NELF-E and NELF-A subunits of the NELF complex are preferentially recruited to DSBs induced upstream transcriptionally active genes in a PARP1-depndent manner to block transcriptional elongation of Pol II (Awwad et al., 2017; Polo, 2017). On the other hand, histone post-translational modifications (PTMs) such as phosphorylation, methylation, ubiquitination, and acetylation are involved in remodeling the chromatin to establish a repressive environment at DSB sites (Cao et al., 2016; Machour and Ayoub, 2020; Sharma and Hendzel, 2019). Of note, we have recently shown that histone lysine crotonylation (Kcr), enriched at transcriptionally active chromatin (Sabari et al., 2015; Tan et al., 2011), is locally and transiently reduced at DNA damage sites, inflicted by ionizing radiation, etoposide and UV radiation, in an HDAC-dependent manner (Abu-Zhayia et al., 2019). However, it remains unknown whether Kcr reduction explicitly occurs at DSB sites and contributes to DSB-induced transcriptional silencing.

Several studies demonstrated that either pharmacological inhibition of ATM, DNA-PK, PARP1 or depletion of silencing factors such as, NELF-E, PBAF and the transcriptional elongation factor, ENL, alleviates transcriptional silencing at DSB sites and impairs HDR and NHEJ repair pathways (Awwad et al., 2017; Kakarougkas et al., 2014; Machour and Ayoub, 2020; Pankotai et al., 2012; Shanbhag et al., 2010; Ui et al., 2015). In this regard, we demonstrated that the chromodomain Y-like (CDYL1) repressor is rapidly recruited to DSBs, in a PARP1-dependent manner, to promote HDR of DSBs (Abu-Zhayia et al., 2018). Interestingly, CDYL1 underpins DSB-induced transcriptional silencing plausibly via recruiting EZH2 methyltransferase to promote H3K27me3 at DSB sites (Abu-Zhayia et al., 2018; Liu et al., 2017b). CDYL1 contains a chromodomain (CD), a central hinge region and an enoyl-CoA hydratase-like (ECH) domain. The CD binds the repressive methyl marks di- and tri-methylated lysine 9 and lysine 27 of histone H3 (H3K9me2/3, H3K27me2/3). The ECH domain is essential for CDYL1 multimerization (Abu-Zhayia et al., 2018; Franz et al., 2009) and mediates its interaction with histone deacetylases 1 and 2 (HDAC1/2) (Escamilla-Del-Arenal et al., 2013). In addition, the region containing the ECH domain regulates CDYL1 recruitment to DNA damage inflicted by laser-microirradiation (Abu-Zhayia et al., 2018). Interestingly, the ECH domain of CDYL1 negatively regulates histone Kcr by acting as a crotonyl-CoA hydratase to convert crotonyl-CoA to β-hydroxybutyryl-CoA (Liu et al., 2017a). Whether CDYL1 crotonyl-CoA hydratase activity underpins a reduction in Kcr at DNA breakage sites remains unknown.

It is widely accepted that DSB-induced transcriptional silencing prevents clashes between repair and transcription factors at DSB sites and precludes the generation of truncated and aberrant transcripts when DSBs occur within a gene body. Also, the formation of repressive chromatin structures at DSB sites restrains the mobility of the broken DNA ends and keeps them in close proximity to ensure timely repair of DSBs (Capozzo et al., 2017; Caron et al., 2019; Gursoy-Yuzugullu et al., 2016; Kakarougkas et al., 2014; Machour and Ayoub, 2020; Purman et al., 2019; Ui et al., 2020; Ui et al., 2015). Despite this, further experimental evidence is pivotal to determine whether DSB-induced transcriptional silencing is indeed prerequisite for intact DSB repair. Herein, we performed systematic profiling of lysine crotonylation, PanKcr, and crotonylated histone residue, H3K9cr, at hundreds of annotated *Asi*SI-induced DSBs across the genome. We showed that PanKcr and H3K9cr levels are reduced at *Asi*SI-induced DSB sites in a CDYL1-dependent manner. Specifically, we demonstrated that CDYL1 crotonyl-CoA hydratase activity downregulates lysine crotonylation at *Asi*SI-induced DSBs leading to transcriptional silencing, at least in part, through the eviction of the Kcr reader ENL from transcriptional start sites (TSS). Next, we exploited CDYL1-S467A mutant that lost its crotonyl-CoA hydratase activity, to study the functional crosstalk between DSB-transcriptional silencing and repair. We observed that the CDYL1-dependent reduction in H3K9cr at DSBs is critical for DSB-induced transcriptional silencing, but unexpectedly has no noticeable effect on HDR integrity. Therefore, our data favor the notion that local DSB-induced transcriptional silencing is dispensable for the integrity of HDR.

## Results

### Systematic identification of a local decrease in histone lysine crotonylation at *Asi*SI-induced DSB sites

DSB-induced transcriptional silencing is partially mediated by alterations in various PTMs including phosphorylation, methylation, ubiquitination and acetylation (Caron et al., 2019; Machour and Ayoub, 2020). Prompted by these observations, we proposed that lysine crotonylation (Kcr), which marks transcriptionally active chromatin, is downregulated at DSB sites to underpin transcriptional silencing. Herein, we sought to systematically determine the levels of Kcr at the local vicinity of DSB sites at defined and native loci throughout the human genome. To achieve this, we used an elegant DSB-Induced via *Asi*SI (DIvA) system (Iacovoni et al., 2010). This system consists of human osteosarcoma cells (U2OS) stably expressing cytoplasmic *Asi*SI restriction enzyme, with an 8bp recognition sequence, fused to oestrogen receptor (ER) hormone-binding domain. Addition of 4-hydroxy tamoxifen (4OHT) to the growth medium drives the migration of *Asi*SI-ER fusion protein into the nucleus leading to the induction of DSB at annotated sites scattered across the human genome (Figure S1A). To validate the induction of DSBs at *Asi*SI recognition sites, untreated and 4OHT-treated DIvA cells were subjected to γH2AX chromatin immunoprecipitation followed by deep sequencing (ChIP-seq). To identify *Asi*SI sites with highest cutting efficiency, we compared γH2AX peaks in 4OHT treated DIvA cells versus untreated cells in a 1Mb window around each of the annotated 1220 *Asi*SI sites across the human genome (Figure 1A). We chose 214 *Asi*SI sites with the highest enrichment of γH2AX and examined its profile surrounding these sites. In agreement with previous findings (Iacovoni et al., 2010; Iannelli et al., 2017), γH2AX enrichment was observed up to 2Mb surrounding the DSB sites, while no peaks were detected in random genomic loci (Figure S1B). Then, we compared the 214-γH2AX enriched *Asi*SI sites with those that were identified by BLESS and BLISS methodologies from two independent studies (Clouaire et al., 2018; Iannelli et al., 2017). Out of the 214-γH2AX enriched *Asi*SI sites, 64 sites were common with *Asi*SI sites identified by BLISS/BLESS assay (Figure 1B) and exhibit strong γH2AX enrichment following DSB induction when compared to uncut *Asi*SI sites (Figure 1C). Consequently, we sought to determine the precise distribution of Kcr around the 64 *Asi*SI sites. Toward this end, untreated and 4OHT-treated cells were subjected to ChIP-seq using PanKcr antibody. Our results show that Kcr levels at 10kb window around *Asi*SI sites were significantly reduced following DSB induction, while negligible changes were observed around uncut *Asi*SI sites (Figure 1D and S2A). Similarly, ChIP-seq for crotonylated lysine 9 of H3 (H3K9cr) shows a significant reduction at *Asi*SI-induced DSBs (Figure 1E and S2B). Interestingly, a prominent reduction in both PanKcr and H3K9cr levels is observed up until 25kb surrounding *Asi*SI sites (Figure S3A-B). However, while H3K9cr levels return to basal levels at 25kb surrounding DSBs, PanKcr downregulation extends up to 5Mb in 4OHT-treated cells compared to untreated cells, suggesting that crotonylation of other residues rather than H3K9cr is also reduced during DSB induction (Figure S3B). Collectively, our data reveal that PanKcr and H3K9cr levels are substantially reduced at *Asi*SI-induced DSB sites.

**Figure 1:**
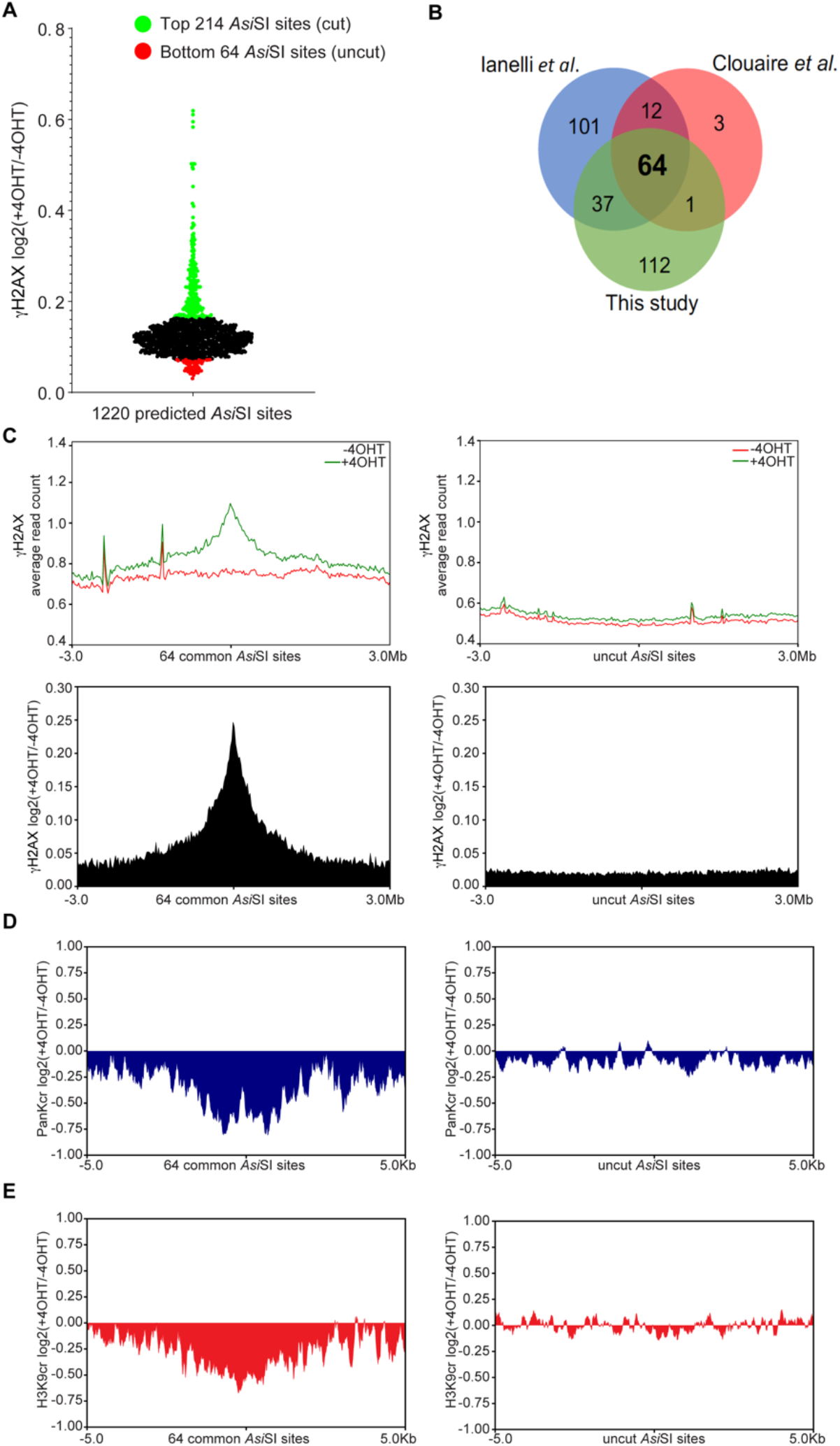
Genome-wide lysine crotonylation profiling revealed local decrease in PanKcr and H3K9cr at *Asi*SI-induced DSBs. **(A)** Dot plot representing ChIP-seq enrichment of γH2AX between 4OHT-treated and untreated U2OS-DIvA cells (expressed as log2 ratio) in a 1Mb window around each of the annotated 1220 *Asi*SI sites of the human genome. 214 *Asi*SI sites with the highest γH2AX enrichment are indicated in green (referred as cut sites) and the 64 *Asi*SI sites with the lowest enrichment are indicated in red (referred as uncut sites). **(B)** Venn diagram plot showing the intersection of the 214 *Asi*SI detected by γH2AX ChIP-seq in this study with previously published cleaved *Asi*SI sites detected by BLESS (Clouaire et al., 2018) or BLISS (Capozzo et al., 2017). **(C)** Average profile of γH2AX enrichment between 4OHT-treated and untreated DIvA cells in 6Mb window surrounding the 64 common cleaved *Asi*SI sites (left) or uncut *Asi*SI sites (right). Top: values are expressed as average normalized CHIP-seq reads. Bottom: values are expressed as average log2 ratio between 4OHT-treated and untreated cells. **(D-E)** Average log2 ratio of PanKcr ChIP-seq (D) or H3K9cr ChIP-seq (E) between 4OHT treated and untreated DIvA cells in 10kb window surrounding the 64-common cut *Asi*SI sites (left) or uncut *Asi*SI sites (right).

### CDYL1 downregulates PanKcr and H3K9cr at *Asi*SI-induced DSBs

Previously, we have shown that CDYL1 is recruited to DSB sites and fosters transcriptional silencing (Abu-Zhayia et al., 2018). In addition, CDYL1 possesses crotonyl-CoA hydratase activity that negatively regulates Kcr (Liu et al., 2017a). On this basis, we hypothesized that CDYL1 attenuates Kcr at DSB sites. To test this hypothesis, we established CDYL1-deficient DIvA cells using shRNA against the 3’UTR region of CDYL1 (Figure 2A). Next, untreated and 4OHT-treated cells were subjected to ChIP-seq analysis using γH2AX, PanKcr and H3K9cr antibodies. We observed that following 4OHT treatment, γH2AX levels are remarkably increased at the 64-*Asi*SI-induced DSBs and spread up to 2Mb (Figure 2B). Notably, CDYL1-deficient cells exhibit higher levels of γH2AX at the 64-*Asi*SI-induced DSBs when compared to control cells (Figure 2C). This finding is in line with our previous data showing that CDYL1-deficient cells exhibit elevated levels of γH2AX after ionizing radiation (Abu-Zhayia et al., 2018). Interestingly, ChIP-seq data of PanKcr in CDYL1-deficient cells revealed a reduction that extended over 5kb surrounding the 64-*Asi*SI-induced DSBs (Figure 2D and Figure S4A). However, the decrease in PanKcr levels in CDYL1-deficient cells is significantly lower than the decrease observed in CDYL1-proficient cells at the 64-*Asi*SI-induced DSBs, suggesting that decrotonylases other than CDYL1 are implicated in regulating Kcr levels at DSB sites (Figure 2E). Unlike PanKcr, no reduction in H3K9cr levels was observed over 5kb surrounding the 64-*Asi*SI-induced DSBs in CDYL1-deficient cells, suggesting that CDYL1 is a master regulator of H3K9cr levels at DSB sites (Figure 2F-G and Figure S4B). Altogether, we concluded that CDYL1 counteracts Kcr levels at DNA breakage sites. To further substantiate the emerging role of CDYL1 in regulating Kcr levels at *Asi*SI-induced DSBs, we focused on three *Asi*SI-sites nearby transcriptionally active genes: MIS12, RBMXL1, and LYRM2, hereafter called DSB-I, DSB-II and DSB-III, respectively. We became interested in these sites because according to ChIP-seq data they exhibit a pronounceable enrichment of γH2AX that correlates with a remarkable reduction in PanKcr and H3K9cr levels, as shown in Figure S5A-C. To confirm the ChIP-seq data, we determined the relative enrichment of γH2AX, PanKcr and H3K9cr using ChIP followed by real-time quantitative PCR (ChIP-qPCR) at DSB-I, DSB-II and DSB-III sites. We observed that following 4OHT treatment, CDYL1-proficient cells display a substantial decrease in PanKcr and H3K9cr levels that inversely correlate with γH2AX levels at DSB-I, DSB-II and DSB-III sites. On the other hand, no reduction in PanKcr and H3K9cr levels was detected in CDYL1-deficient cells, except of a moderate reduction in in PanKcr at DSB-I (Figure 3A-C). These findings are concordant with the aforementioned PanKcr and H3K9cr ChIP-seq data (Figure 2E, G). Next, we sought to determine whether the decrease in Kcr levels is accompanied with CDYL1 accumulation at DSB sites. Intriguingly, ChIP-qPCR for CDYL1 showed a remarkable accumulation of CDYL1 at DSB-I, DSB-II and DSB-III sites following 4OHT treatment (Figure 3D). We concluded therefore that CDYL1 is recruited to DSB sites to counteract Kcr.

**Figure 2:**
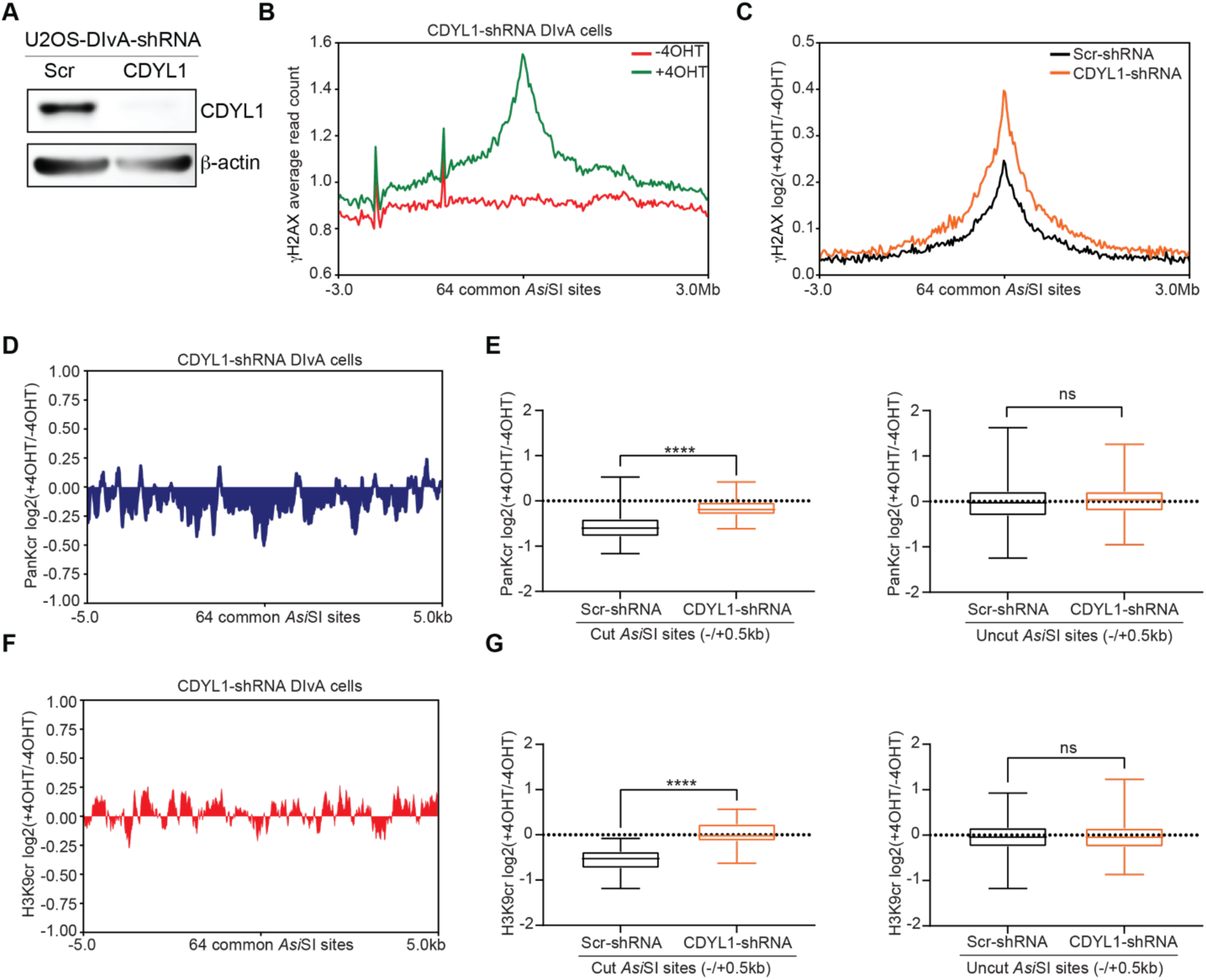
CDYL1 impedes PanKcr and H3K9cr at *Asi*SI-induced DSBs. **(A)** Western blot shows CDYL1 knockdown using shRNA against 3’UTR region of CDYL in DIvA cells. β-actin is used as a loading marker. **(B)** Average normalized ChIP-seq reads of γH2AX between 4OHT-treated and untreated CDYL1-depleted cells in 6Mb window surrounding the 64 common cleaved *Asi*SI sites. **(C)** Average log2 ratio of γH2AX enrichment between 4OHT-treated and untreated DIvA cells in 6Mb window surrounding the 64 common cleaved *Asi*SI sites in DIvA cells expressing scramble or CDYL1-shRNA **(D)** Average log2 ratio of PanKcr ChIP-seq between 4OHT treated and untreated CDYL1-depleted DIvA cells in 10kb window surrounding the 64 common cleaved *Asi*SI sites. **(E)** Boxplots representing log2 ratio of PanKcr between treated and untreated DIvA cells expressing either scramble shRNA or shRNA against CDYL1 in 1kb window surrounding the 64 common cleaved *Asi*SI sites (left) or uncut *Asi*SI sites (right). p-values were calculated using non-parametric Mann-Whitney test. *p<0.01; **p<0.001; ***p<0.0001; ****p<0.00001; ns: not significant. **(F)** as in (D) except H3K9cr average log2 ratio is represented. **(G)** as in (E) except H3K9cr is represented.

**Figure 3:**
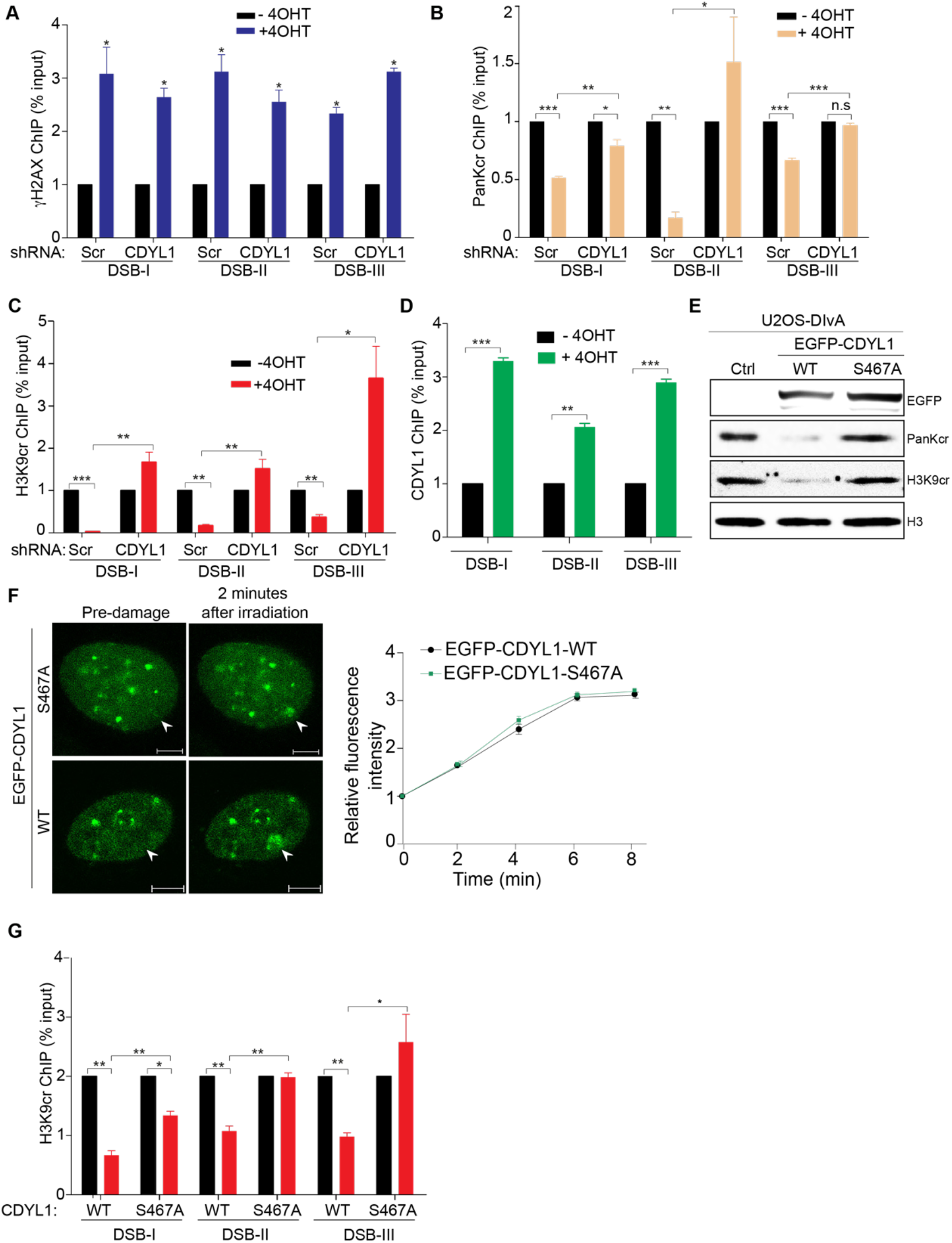
CDYL1 crotonyl-CoA hydratase activity downregulates PanKcr and H3K9cr at *Asi*SI-induced DSBs. **(A-C)** ChIP-qPCR for γH2AX (A), PanKcr (B) and H3K9cr (C) in DIvA cells expressing either scramble or CDYL1 shRNA before and after addition of 300nM 4OHT for 4hrs. Data represented as % of input. The mean of normalized ChIP efficiency and SEM of three representative experiments are shown (out of 3 independent repeats). **(D)** CDYL1 is recruited to *Asi*SI-induced DSBs. ChIP-qPCR was performed using antibody against CDYL1 in DIvA cells that were left untreated or treated for 4hrs with 300nM 4OHT. Data represented as % of input. **(E)** Western blot analysis showing that expression of CDYL1-WT but not CDYL1-S467A leads to a decrease in PanKcr and H3K9cr levels. H3 is used as a loading marker. **(F)** Laser-microirradiation shows a comparable recruitment of EGFP-CDYL1-WT and EGFP-CDYL1-S467A mutant to DNA breakage sites (marked with white arrowheads) in DIvA cells. Graph in the right displays fold increase in the relative fluorescence intensity of EGFP-CDYL1-WT and EGFP-CDYL1-S467A at damaged sites. **(G)** as in C, except that ChIP was performed in CDYL1-deficient DIvA cells that have been complemented with vectors expressing either CDYL1-WT or CDYL1-S467A mutant. The mean of normalized ChIP efficiency and SEM of three representative experiments are shown (out of 3 independent repeats). Error bars represent standard deviations from the mean. *p<0.01; **p<0.001; ***p<0.0001.

### CDYL1 crotonyl-CoA hydratase activity counteracts PanKcr and H3K9cr at DSB sites

It was shown that CDYL1 crotonyl-CoA hydratase activity negatively regulates Kcr during spermatogenesis (Liu et al., 2017a). Therefore, we assumed that CDYL1 attenuates Kcr at DSB sites via its crotonyl-CoA hydratase activity. To test this assumption, we took advantage of a CDYL1-S467A mutant that lacks its crotonyl-CoA hydratase activity (Liu et al., 2017a). First, we validated that this mutant indeed lost its crotonyl-CoA hydratase activity. Western blot analysis shows that overexpression of CDYL1-WT reduces PanKcr and H3K9cr levels, while overexpression CDYL1-S467A has no detectable effect on Kcr levels, suggesting that CDYL1-S467A mutant indeed lost its crotonyl-CoA hydratase activity (Figure 3E). Since S467 residue is located within the ECH domain that is known to regulate CDYL1 recruitment to DNA damage sites (Abu-Zhayia et al., 2018), we sought to determine whether S467A mutation affects CDYL1 recruitment to DNA lesions. Laser microirradiation assay revealed comparable recruitment kinetics of CDYL1-S467A and CDYL1-WT to DNA breakage sites, suggesting that S467A mutation has no detectable effect on CDYL1 recruitment to DNA damage sites (Figure 3F). To study the role of CDYL1 crotonyl-CoA hydratase activity on DSB-induced reduction in H3K9cr, we expressed either CDYL1-WT or CDYL1-S467A mutant in CDYL1-deficient DIvA cells expressing shRNA sequence that targets the 3’UTR region of CDYL1. Complementing CDYL1-deficient cells with CDYL1-WT leads to a reduction in H3K9cr at *Asi*SI-induced DSB-I, DSB-II, DSB-III sites as seen by H3K9cr ChIP-qPCR analysis. By contrast, expressing CDYL1-S467A mutant fails to elicit reduction in H3K9cr levels at these DSB sites (Figure 3G). Altogether, our data demonstrate that CDYL1 counteracts Kcr at DSB sites via its crotonyl-CoA hydratase activity.

### Reduction in histone lysine crotonylation correlates with DSB-induced transcriptional silencing

In an attempt to understand the functional relevance of Kcr reduction following DSB induction, we analyzed the genome-wide distribution of PanKcr and H3K9cr in U2OS-DIvA cells. In agreement with previous reports (Gowans et al., 2019; Li et al., 2016; Liu et al., 2018; Tan et al., 2011), PanKcr signal was predominantly enriched at TSS of active genes (Figure S6A). Moreover, we detected similar enrichment of H3K9cr at TSSs suggesting that H3K9cr is also associated with active transcription (Figure S6B). Consequently, our findings postulate that the previously reported transcriptional silencing at *Asi*SI sites (Iannelli et al., 2017) correlates with a reduction in H3K9cr. To substantiate this correlation, we looked at H3K9cr profile at *Asi*SI sites within TSS of genes that exhibit a strong reduction in RNA transcription (Iannelli et al., 2017) (Figure S6C). Intriguingly, the reduction in H3K9cr is significantly more prominent at these sites compared to other *Asi*SI sites that don’t exhibit pronounced reduction in transcription activity (Figure S6C). In addition, DSB-induced reduction in PanKcr and H3K9cr correlates with a previously reported decrease in H3K79me2 (Clouaire et al., 2018), a known modification enriched at transcriptionally active genes (Steger et al., 2008) whose decrease contributes to transcriptional shutdown (Godfrey et al., 2019) (Figure S6D). Noteworthy, we did not find any correlation between H3K9cr and the histone modification H3K9me3, which was shown to remain unchanged at *Asi*SI-induced DSBs (Clouaire et al., 2018) (Figure S6D). This finding supports the notion that the reduction in H3K9cr, rather than increase in H3K9me3, is critical for downregulating transcription at *Asi*SI-induced DSBs.

### The CDYL1-dependent decrease in Kcr underpins DSB-induced transcriptional silencing and promotes ENL eviction from TSS nearby DSB sites

The impact of CDYL1 crotonyl-CoA hydratase activity on DSB-induced transcriptional silencing is unknown. To explore this, we monitored the expression of MIS12 and RBMXL1 genes that are adjacent to DSB-I and DSB-II sites, respectively. In agreement with a previous report (Iannelli et al., 2017), 4OHT treatment induces transcriptional silencing of MIS12 and RBMXL1 genes in control DIvA cells. Intriguingly, however, we revealed that CDYL1 depletion alleviates DSB-induced silencing of MIS12 and RBMXL1 (Figure 4A-B). Moreover, complementing CDYL1-deficient DIvA cells with a vector expressing CDYL1-WT restores DSB-induced transcriptional silencing of MIS12 and RBMXL1. In contrast, complementing CDYL1-deficient DIvA cells with CDYL1-S467A mutant fails to restore DSB-induced silencing (Figure 4A-B), which is in concordant with the lack of Kcr reduction at DSB sites observed in Figure 3G. Altogether, our data provide firm evidence that CDYL1 crotonyl-CoA hydratase activity mediates the reduction in Kcr at DSB sites to ensure transcriptional silencing. Concordantly, pre-treating DIvA cells with sodium crotonate, which leads to an extreme increase in histone Kcr levels (Sabari et al., 2015) (Figure S7A), not only precludes DSB-induced transcriptional silencing of RBMXL1 gene but also enhances its transcription level (Figure S7B). Next, we sought to shed mechanistic insights into how CDYL1-dependent downregulation of Kcr promotes DSB-induced transcriptional silencing. We became interested in the transcriptional elongation factor, ENL, because it is known as a Kcr reader (Li et al., 2016), and its displacement from chromatin reduces RNA Pol II initiation and elongation (Erb et al., 2017; Wan et al., 2017). On this basis, we proposed that CDYL1-dependent downregulation of Kcr triggers ENL displacement from TSS located nearby *Asi*SI-induced DSBs to underpin DSB-induced silencing. To test this, we performed ChIP-qPCR for ENL in CDYL1-proficient and -deficient DIvA cells. Our results show that DSB induction reduces ENL levels at MIS12 and RBMXL1 TSSs, presumably due to the decrease in Kcr levels at DSB-I and DSB-II. In support of this, CDYL1 depletion, which blocks Kcr decrease at DSB sites (Figure 3B, C), eminently reduces ENL dispersal from DSB-I and DSB-II (Figure 4C). This finding suggests that CDYL1 together with other yet unknown factors contribute to ENL displacement from TSS following DSB induction. Altogether, we identify a novel pathway to ensure DSB-induced transcriptional silencing via CDYL1-dependent eviction of ENL from TSSs and hence revealed a previously unrecognized crosstalk between CDYL1 and ENL to foster DSB-induced silencing.

**Figure 4:**
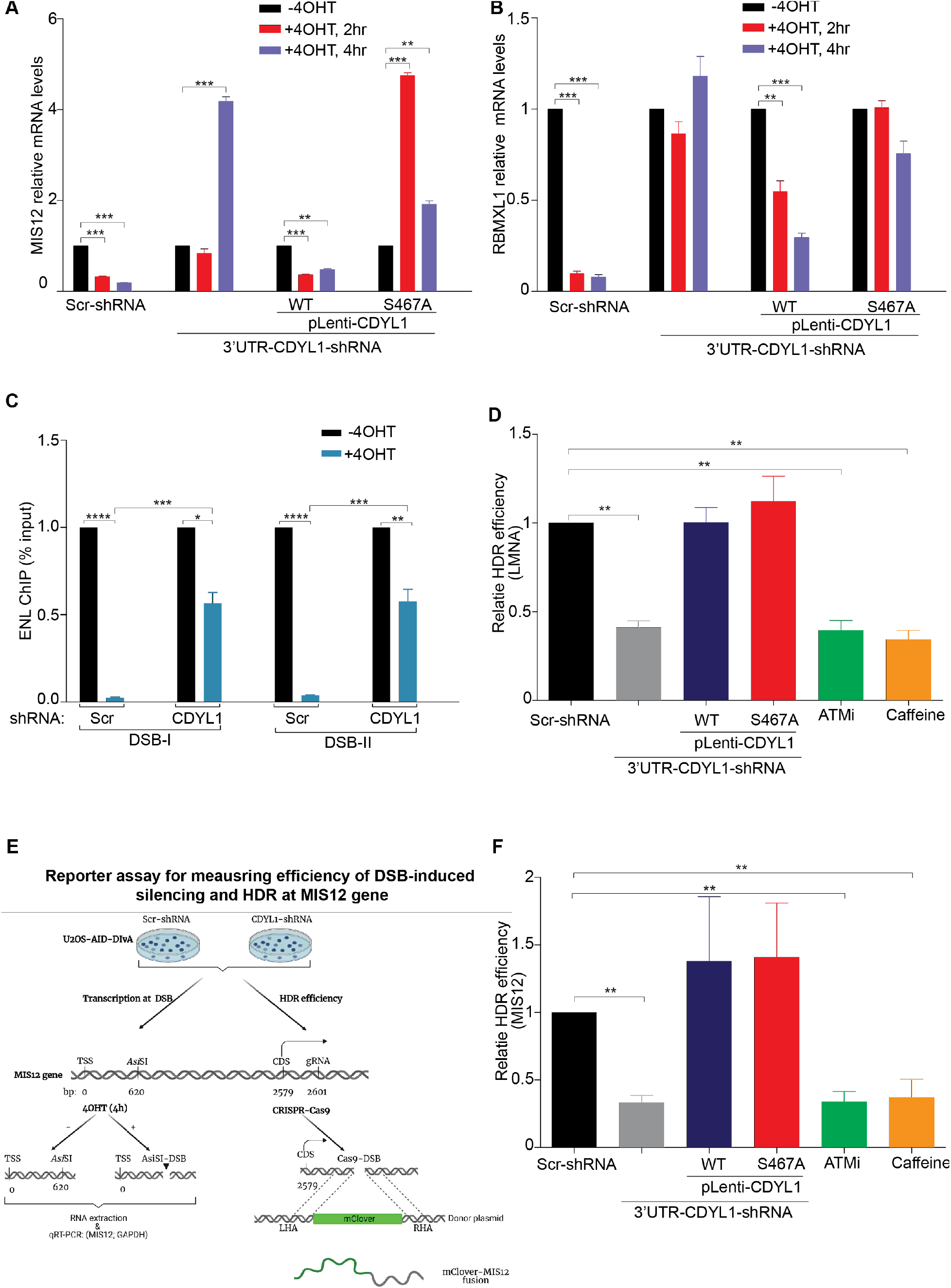
CDYL1 crotonyl-CoA hydratase activity underpins DSB-induced transcriptional silencing but dispensable for HDR. **The CDYL1-dependent decrease in Kcr is dispensable for HDR and triggers ENL eviction to ensure DSB-induced transcriptional silencing. (A-B)** CDYL1 crotonyl-CoA hydratase activity fosters DSB-induced transcriptional silencing of MIS12 and RBMXL1 genes. mRNA expression levels of MIS12 (A) and RBMXL1 (B) were measured before or after 4OHT treatment for 4hrs in CDYL1-proficient and -deficient DIvA cells, and in CDYL1-deficient DIvA cells expressing either CDYL1-WT or CDYL1-S467A mutant. Gene expression was normalized to the levels of GAPDH transcript. *p<0.01, **p < 0.001, ***p < 0.0001. **(C)** CDYL1 activity promotes ENL removal from TSS of MIS12 and RBMXL1 genes. ChIP-qPCR for ENL in DIvA cells expressing either scramble or CDYL1 shRNA before and after 4OHT treatment. Data represented as % of input. The mean of normalized ChIP efficiency and SEM of three representative experiments are shown (out of 3 independent repeats). **(D)** CDYL1 depletion impairs HDR of endogenous DSBs induced by Cas9 endonuclease upstream the LMNA gene. CDYL1-profcient and deficient DIvA cells were transfected with fluorescence (Clover)-based reporter and HDR efficiency was determined at 72hrs after transfection, as described in materials and methods. Results shown are typical of 3 independent experiments. Data are presented as mean ± SD of three independent experiments. *p<0.01, **p < 0.001, ***p < 0.0001, ns: not significant. (**E**) Schematic representation of cell-based reporter assay for measuring both DSB-induced transcriptional silencing and HDR efficacy at MIS12 gene. (**F**) CDYL1 hydratase activity promotes is dispensable for HDR of Cas9-induced DSB at MIS12 gene. HDR efficiency was determined as in (D).

### DSB-induced transcriptional silencing is dispensable for HDR

Herein, we sought to revisit a fundamental belief concerning the essentiality of DSB-induced transcriptional silencing for the integrity of DSB repair, particularly HDR. Given that CDYL1 promotes both DSB-induced silencing and HDR (Abu-Zhayia et al., 2018), and that CDYL1 crotonyl-CoA hydratase activity is critical for DSB-induced silencing, we planned to determine the effect of CDYL1 hydratase activity on HDR. To achieve this, we tested HDR integrity in CDYL1-deficient DIvA cells complemented with vectors expressing either CDYL1-WT or CDYL1-S467A mutant. We used an established fluorescence (Clover)-based reporter that measures HDR efficiency of Cas9-induced DSB at the LMNA gene (Pinder et al., 2015). Surprisingly, although CDYL1-S467A expression didn’t restore DSB-induced silencing, it rescues the defective HDR of Cas9-induced DSB at the LMNA gene to a similar extent as CDYL1-WT (Figure 4D). This unanticipated result demonstrates that CDYL1 crotonyl-CoA hydratase activity is critical for DSB-induced silencing but not for intact HDR, suggesting that DSB-induced silencing is not prerequisite for proper HDR of DSBs. To further substantiate this finding, we sought to determine the effect of CDYL1 crotonyl-CoA hydratase activity on DSB-induced transcriptional silencing and HDR at the same genomic locus nearby MIS12 gene. The rationale behind testing HDR integrity at the MIS12 is that, as we showed, CDYL1 accumulates at DSB-I nearby MIS12 gene, downregulates Kcr and fosters its transcriptional silencing (Figure 3C, D and Figure 4A). Toward this end, we established a Clover-based reporter designated to measure HDR of *Asi*SI-induced DSB at the MIS12 gene (Figure 4E). Intriguingly, we observed that while expression of CDYL1-S467A mutant in CDYL1-deficient DIvA cells fails to underpin DSB-induced silencing, it restores the integrity of HDR of *Asi*SI-induced DSB at the MIS12 gene (Figure 4F). Collectively, these findings functionally uncouple the roles of CDYL1 in HDR and DSB-induced transcriptional silencing and suggest that they may occur independently.

## Discussion

In this study, we systematically mapped histone Kcr, a known modification associated with active transcription (Kollenstart et al., 2019; Li et al., 2016; Sabari et al., 2015; Tan et al., 2011; Wan et al., 2019), before and after DSB induction. Our data uncover a novel role of CDYL1 crotonyl-CoA hydratase activity in downregulating PanKcr and H3K9cr at DSB sites that triggers ENL eviction from TSS site to ensure transcriptional silencing. In addition, our data functionally uncouples the dual role of CDYL1 in fostering transcriptional silencing and HDR of DSBs. Unexpectedly, we showed that alleviation of DSB-induced transcriptional silencing has no discernable effect on HDR, suggesting that DSB-transcriptional silencing and HDR may occur independently (Figure S8). Future studies will be required to address whether DSB-induced transcriptional silencing is required for the integrity of other DSB repair pathways, such as NHEJ. Whilst several studies implicated histone PTMs such as phosphorylation methylation, ubiquitination and acetylation in DSB-induced silencing (Caron et al., 2019; Machour and Ayoub, 2020), here we identified histone Kcr as a novel and critical player in regulating transcriptional silencing at DSBs. Future works will be required to shed light onto the crosstalk and the functional interdependencies between the different PTMs to underpin DSB-induced transcriptional silencing. In this regard, we and other have shown that CDYL1 promotes EZH2 methyltransferase recruitment to DSB sites and consequently facilitates H3K27me3 repressive mark (Abu-Zhayia et al., 2018; Liu et al., 2017b). Therefore, we propose that CDYL1 fosters DSB-induced silencing through downregulating and upregulating Kcr and H3K27me3, respectively. The interdependence between Kcr downregulation and H3K27me3 at DSB sites and their contribution to silencing deserves future research. Also, the spatiotemporal crosstalk between the different crotonylase and decrotonylase enzymes at DSB sites remains unknown and needs rigorous investigation. For example, in addition to CDYL1, we have recently shown that HDACs downregulate Kcr following DNA damage (Abu-Zhayia et al., 2019). This raises a question of whether simultaneous inhibition of HDAC and CDYL1 has a synergistic effect on DSB-induced transcriptional silencing and repair. Also, what is the role, if any, of other known decrotynalses in modulating Kcr levels at DSB sites?

In this study, we demonstrated that the levels of both PanKcr and H3K9cr decrease at *Asi*SI-induced DSBs which is consistent with our previous findings showing that Kcr level is transiently reduced following DNA damage inflicted by ionizing radiation, UV, and Etoposide treatment (Abu-Zhayia et al., 2019). Notably, while the decrease in H3K9cr is up to 25kb, the decrease in PanKcr spread over 5Mb surrounding *Asi*SI-induced DSB (Figure S3). This indicates that the crotonylation of histone lysine residues other than H3K9cr are also reduced in response to DSB induction. Moreover, since Kcr has been identified in both histone and non-histone proteins (Wu et al., 2017; Xu et al., 2017; Yu et al., 2020), it would be exciting to determine whether the crotonylation of nonhistone proteins (*e*.*g*., DNA repair proteins) is altered during DNA damage in a CDYL1-dependent manner. An additional question that stems from this work is whether histone Kcr decrease following DSB induction is influenced by the cell cycle stage and/or chromatin structure surrounding DSB sites. In addition, since we have shown that the reduction in histone Kcr occurs following different types of DNA lesions including UV damage (Abu-Zhayia et al., 2019), it will be required to determine whether CDYL1 crotonyl-CoA hydratase activity regulates Kcr levels following various types of DNA lesions other than DSBs.

How does CDYL1 regulate transcription activity at DSB sites? Previously, we and others have shown that CDYL1 is recruited in a PARP1-dependent manner and recruits EZH2 to foster trimethylation of H3K27 (Abu-Zhayia et al., 2018; Liu et al., 2017b). Herein, we identified an additional pathway that involves CDYL1-dependent reduction of H3K9cr at DSB sites, which subsequently leads to the removal of the Kcr reader ENL, a known transcription elongation factor (Li et al., 2016; Luo et al., 2012; Schulze et al., 2009; Zhou et al., 2012), from the TSS of MIS12 and RBMXL1 genes. Our work highlights therefore a previously unrecognized crosstalk between CDYL1 and ENL to ensure DSB-induced silencing. In this regard, it was shown that after DSB induction ATM phosphorylates ENL, which triggers the recruitment of PRC1 complex to ubiquitinate H2A and induce transcriptional silencing at DSB sites (Ui et al., 2015). Taken together, we propose that ENL exerts its function in DSB-induced silencing through two distinct mechanisms that involve its eviction from TSS of active genes and recruitment of repressive factors to the transcriptional elongation sites. Future work is needed to determine whether the reduction of Kcr at DSB sites leads to the eviction of other known histone Kcr readers such as Yaf9, Taf14, AF9 and Sas5 (Andrews et al., 2016; Li et al., 2016; Zhang et al., 2016; Zhao et al., 2016) and if so, to determine their role in DSB-induced silencing. In a broader context, our data uncover a potential synergism between ATM that phosphorylates ENL, and PARP1 that promotes CDYL1 recruitment to DSB sites, to foster DSB-induced silencing.

While DSB-induced silencing is critical to avoid transcribing broken genes, its impact on DSB repair is not fully understood. A leading dogma is that the repressive chromatin organization immobilize the DSB ends and keep them close to each other to foster timely DSB repair (Capozzo et al., 2017; Caron et al., 2019; Gursoy-Yuzugullu et al., 2016; Kakarougkas et al., 2014; Machour and Ayoub, 2020; Purman et al., 2019; Ui et al., 2020; Ui et al., 2015). This dogma is supported by at least two independent observations showing that the alleviation of DSB-induced transcriptional silencing following the depletion of either the chromatin remodeling PBAF complex or DYRK1B kinase leads to defective DSB repair. Moreover, they demonstrated that global inhibition of transcription activity by DRB restored the integrity of DSB repair measured by NHEJ and comet assays (Dong et al., 2020; Kakarougkas et al., 2014). It remains unknown however the impact of DSB-induced silencing on HDR integrity. To study the crosstalk between DSB-induced silencing and HDR, we established an elegant system that enables us to measure transcription levels and HDR efficiency at the same locus (MIS12 gene). Then, we took advantage of CDYL1-S467A mutant that lost its DSB-induced silencing activity but retains the ability to promote HDR. Complementing CDYL1-deficient cells with either wild-type or CDYL1-S467A mutant allowed us to functionally uncouple the silencing and the repair activity of CDYL1 (Figure 4). Indeed, induction of DSB at *Asi*SI site within the MIS12 gene in CDYL1-deficient cells expressing CDYL1-S467A mutant failed to induce transcriptional silencing of MIS12 but restored HDR efficiency. Our study therefore provides firm evidence that DSB-induced silencing is not prerequisite for intact HDR. These apparently contradictory observations, concerning the essentiality of transcriptional silencing to intact DSB repair, could be attributed to the different tested DSB repair pathways. Altogether, we postulate that while DSB-induced silencing is required for NHEJ, it is dispensable for HDR. Alternatively, the rescue of DSB repair in DYRK1B and PBAF-deficient cells may arise from indirect and undesirable side effects of the spurious transcription inhibition using DRB.

## Acknowledgments

We are grateful to Gaëlle Legube for providing us with the powerful U2OS-DIvA system. We thank Jing Liang (Peking University) and our lab members for critical reading of the manuscript, Graham Dellaire for providing the pCR2.1-CloverPML and pX330-LMNA-gRNA plasmids for the homologous recombination assay. Research in the Ayoub lab is supported by grants from the Israel Science Foundation (2511/19), ISF-NSFC fund (# 2511/18), Israel Cancer association (20200080). E.R.A. is supported by Clore fellowship. F.E.M. is supported by Irwin and Joan Jacob fellowship. L.A.B is supported by the VATAT fellowship for outstanding minority MSc students. N.A. is supported by the Neubauer Family foundation.

## Author Contributions

E.R.A, F.E.M. and L.A.B helped in writing the manuscript and wrote the materials and methods. E.R.A performed the ChIP-seq experiments described in figure 1, 2, and supplementary figures S1-S6 and prepared S8. Also, she performed the experiments described in figures 2A, 3A-G, 5A-B, D, E-F and S7. F.E.M developed bioinformatic analysis pipelines, analyzed all ChIP-seq data, prepared figures 1-2, supplementary Figures S1, S2, S3, S4, S5, and S6 and participated in conceiving the study. L.A.B established and calibrated the DSB-Induced via *Asi*SI (DIvA) system in our lab and prepared some of the constructs and the cell lines used in in this manuscript. She performed the experiment described in figure 4C and did some of the biological repeats of the ChIP-seq and ChIP-qPCR assays. B.B.O helped in performing the experiments designed to establish CDYL1 role in counteracting Kcr at DSB sites and in editing the manuscript. N.A. conceived the study, planned the experiments, analyzed the data and wrote the original draft.

## Competing Interests

The authors declare that they have no competing interest.

## Materials and methods

### Cell lines and treatments

U2OS-DIvA (AID-*Asi*SI-ER-U20S) cells were cultured in Dulbecco’s modified Eagle’s medium (DMEM) supplemented with 10 % heat-inactivated fetal bovine serum (Gibco), 2mM L-glutamine (Gibco), 100unit/mL penicillin, 100μg/mL streptomycin (Gibco) and 1mM Sodium Pyruvate in the presence of 400μg/mL G418 at 37°C and 5% CO2. U2OS-DIvA cells infected with lentiviral plasmids expressing either scramble or CDYL-shRNA sequence and were cultured in the presence of 1 μg/mL puromycin. For AsiSI-dependent DSB induction, cells were treated with 300nM (z)-4-Hydroxytamoxifen (4OHT) for the indicated time points. Cell transfections with plasmid DNA were performed using Polyethylenimine (PEI) following the manufacturer’s instructions. Sodium crotonate was prepared by dissolving solid crotonic acid in PBSX1 followed by titration with sodium hydroxide to pH 7.4

### Generation of lentiviral particles and cell transduction

Gene expression knockdown using shRNA sequences was performed as previously described (Khoury-Haddad et al., 2014). First, scramble short hairpin oligonucleotides and short hairpin oligonucleotide directed against CDYL1 were annealed and inserted into pLKO.1-TRC lentiviral vector digested with *Eco*RI and *Age*I. The newly generated lentiviral vectors were validated by DNA sequencing. For the rescue experiments, wild type CDYL1 gene or S467A mutant were cloned into pCSC-SP-PW-(GENE)-IRES/GFP lentiviral plasmid as listed in Table 2. To generate viral particles containing the indicated constructs, HEK293T cells plated in 10cm dish were co-transfected with the lentiviral plasmid (1.64pmol), together with viral packaging plasmids; psPAX2 (1.3pmol) and pMD2.G (0.72pmol). Media containing the viral particles were collected 48 h post-transfection and filtered with 0.45μm filters. Viral particles were used to infect the DIvA cells. Cells infected with shRNA expressing plasmids were grown in the presence of 1μg/ml Puromycin.

### Western blot

Protein extracts were prepared using Hot-lysis buffer (1% SDS, 5mM EDTA, 50mM Tris, pH 7.5) as detailed in (Khoury-Haddad et al., 2014). Antibodies used for western blot and their dilutions are described in Table 1.

### Laser micro-irradiation

Cells were subjected to laser microirradiation as previously described (Khoury-Haddad et al., 2014). Briefly, cells were plated on flourodish (Ibidi; Cat#81158) and pre-sensitized with 1μg/μl Hoechst 3334 dye for 10 min at 37°C. Laser microirradiation was executed using an LSM-700 inverted confocal microscope equipped with CO2 module and 37°C heating chamber. DNA damage was induced by micro-irradiating a single region in the nucleus with 15 iterations of 405 nm laser beam. Time-laps images were acquired using 488 nm laser. Signal intensity at damaged sites was measured using Zen 2009 software.

### RNA Isolation, reverse transcription, and quantitative real-time PCR

Total RNA was isolated from cells using the TRIzol reagent according to the manufacturer’s instructions (Ambion). RNA was treated with DNase (Ambion) for 20 min to remove any potential residual genomic DNA contamination. 1 μg RNA was used for cDNA synthesis using qScript cDNA Synthesis Kit (Quanta) with random primers. mRNA levels were measured using Step-One-Plus real-time PCR System (Applied Biosystems) against the indicated primers and the Fast SYBR Green Master mix (Applied Biosystems). Data analysis and quantification were performed using StepOne software V2.2 supplied by Applied Biosystems. GAPDH gene was used as a housekeeping gene.

### Endogenous homologous recombination assay

Homologous recombination repair assay was performed using Cas9-mediated knock-in of green-fluorescent mClover into the first exon of the LMNA gene, as previously described (Pinder et al., 2015). Briefly, cells were plated in 6-well plates and co-transfected with 1.6μg pX330-LMNA-gRNA1 plasmid expressing Cas9 and gRNA targeting exon 1 of LMNA gene and 0.4μg pCR2.1-CloverLamin plasmid containing HDR donor sequence. Also, 0.4 μg pDsRed-Monomer-C1 was included per transfection as a transfection control unless otherwise indicated in the figure legend. 12−16h post transfection, culture medium was renewed and where indicated, ATMi (5μM) or Caffeine (4mM) were added. 72 h post-transfection, cells were collected and analyzed by flow cytometry. HDR efficiency was calculated as %mClover expressing cells out of DsRed-Monomer positive cells. To test HDR integrity nearby MIS12 gene, we generated pX330-MIS12-gRNA1 plasmid containing Cas9 and gRNA targeting MIS12 gene and pCR2.1-Clover-MIS12-Donor plasmid containing HDR donor sequence (as described in table 2) and proceeded as described above.

### Cell cycle analysis by flow cytometry

Flow cytometric analysis was performed as previously described (Abu-Zhayia et al., 2018). Briefly, cells were fixed with ice-cold 75% ethanol. DNA was stained with 100 mg/ml propidium iodide (Sigma-Aldrich) in phosphate buffer solution (PBS) containing 0.5mg/ml DNase free RNase A (Sigma-Aldrich) and 0.1% Triton-X-100. Samples were analyzed using a BD LSR-II flow cytometer (Becton Dickinson). Data were analyzed with FCS express software.

### Chromatin immunoprecipitation

ChIP assays were carried out as previously described (Aymard et al., 2014). Briefly, cells were plated in 150-mm dishes. When reached ∼80% confluency, cells were treated with 300nM 4OH for 4 hrs. Cells were fixed with 1% PFA for 10 minutes at room temperature, and cross-linking was stopped with 0.125 M Glycine for 5 min. After cell lysis, DNA was sheared to the size of 300– 500 bp using a Vibra cell sonicator (15 sec ON, 30 sec OFF, 35% duty, 20 cycles). Five percent of each supernatant was used as input control and processed with the cross-linking reversal step. Chromatin was immunoprecipitated using 2ug of the indicated antibodies and Protein A/G magnetic beads. Following reverse cross-linking; the precipitated DNA was purified using the Macherey-Nagel Nucleospin kit. Both input and IP samples were analyzed by real-time PCR using the primers listed in table 3. ChIP efficiency was normalized to the total percent of input DNA immunoprecipitated. For ChIP-seq, ∼5 ng of purified DNA (average size 300-500 bp) was used for library preparation and sequenced using illumina NovaSeq S1 (single-end, 100bp reads) at The Medicinal Chemistry institute of the Nancy and Stephen Grand Israel National Center for Personalized Medicine, Weizmann Institute of Science. Average read depth of 24 million reads per sample was obtained. Quality assessment was performed using FastQC to verify the sequencing data pass the quality standard (http://www.bioinformatics.babraham.ac.uk/projects/fastqc/).

Publicly available previously reported high-throughput sequencing data (Clouaire et al., 2018) for H3K9me3 and H3K79me2 CHIP-seq in DIvA cells (-/+4OHT) were downloaded from ArrayExpress: E-MTAB-5817 and re-analyzed using the pipeline described in this study.

### ChIP-seq read processing

All samples were aligned to the GRCh38/hg38 human genome assembly using bowtie2(Langmead and Salzberg, 2012). Aligned reads in SAM format were then converted to BAM format and sorted, indexed, and quality filtered using SAMtools (Li et al., 2009). Next, quality control and GC-bias correction were performed using a suite of python tools from deepTools (Ramirez et al., 2016). For each aligned dataset in BAM format, bamCoverage (deepTools) tool was used to compute the coverage and normalize each sample using Reads Per Kilobase per Million mapped reads. Coverage files for each biological replicate in bigwig format were combined using bigwigCompare “add” tools (deepTools) and exported for further processing.

### Averaged ChIP-seq profiles

Averaged ChIP-seq profiles around specific AsiSI and non-AsiSI sites were computed from coverage tracks using computeMatrix (deepTools) at each 10bp bin, except for larger windows (1 Mb scale) where data were smoothed using 25 kb bins. Log2 ratio tracks were generated using the bigwigCompare tool from deepTools (http://deeptools.readthedocs.io) bigwig tracks -/+4OHT as inputs and average log2 ratio around specific AsiSI and non-AsiSI sites were computed using computeMatrix as described above. Normalized read count profiles and log2 ratio profiles were plotted using plotProfile (deepTools).

### In silico prediction of *Asi*SI sites

Genomic coordinates containing exact match of *Asi*SI restriction sites (5′-GCGATCGC-3′) were extracted from GRCh38/hg38 genome assembly using ad-hoc R script.

### Generation of random non-*Asi*SI sites

A control set of 64 random non-*Asi*SI sites in the genome was generated using bedTools “random” (Quinlan and Hall, 2010) by computing 128 random position on the entire genome (GRCh38/hg38) and then randomly choosing 64 sites that meet the following criteria: not on chromosome Y and at least 1MB away from the top 214 cleaved *Asi*SI sites according to γH2AX CHIP-seq data generated in this study (determined by bedTools “intersect”).

### Determination of the top *Asi*SI sites

To identify AsiSI sites with highest cutting efficiency, average log2 ratio of γH2AX CHIP-seq before and after 4OHT treatment was computed in 1MB window around the 1220 predicted AsiSI sites. 300 *Asi*SI sites with the highest log2 ratio in U2OS-DIvA-Scramble-shRNA and U2OS-DIvA-CDYL1-shRNA cells were overlapped, generating a set of 214 *Asi*SI sites with the highest γH2AX. Next, these 214 AsiSI sites were compared with previously published BLISS data (Clouaire et al., 2018; Iannelli et al., 2017) and the common 64 sites between the 3 studies were extracted. 64 *Asi*SI sites with the lowest log2 ratio in both U2OS-DIvA-Scramble-shRNA and U2OS-DIvA-CDYL1-shRNA cells were subsequently extracted and used as control uncut *Asi*SI sites. All overlaps were computed using bedTools “intersect”.

### Descriptive statistics

Average log2 ratio between untreated and 4OHT treated samples was calculated over the specified window around AsiSI and non-*Asi*SI sites and represented in a boxplot. The box ends represent the first and third quartiles, the center line represents the median, and the whiskers represent the minimum and maximum values. Nonparametric unpaired Mann-Whitney-Wilcoxon test was performed to test the significance of the differences between the samples. Spearman correlation coefficients were calculated to examine the correlation between different samples and treatments.

## Statistical analysis

All statistical representations and analysis were performed using Graphpad Prism 9.0.

## Data availability

The accession number for the high-throughput sequencing data reported in this paper is ArrayExpress: E-MTAB-10550.

## Notes

### Competing Interest Statement

The authors have declared no competing interest.

