## Supplemental Figures and methods for "Systematic identification of a CDYL1-dependent decrease in lysine crotonylation at DNA double-strand break sites functionally uncouples transcriptional silencing and repair"

**Supplementary Figures:**

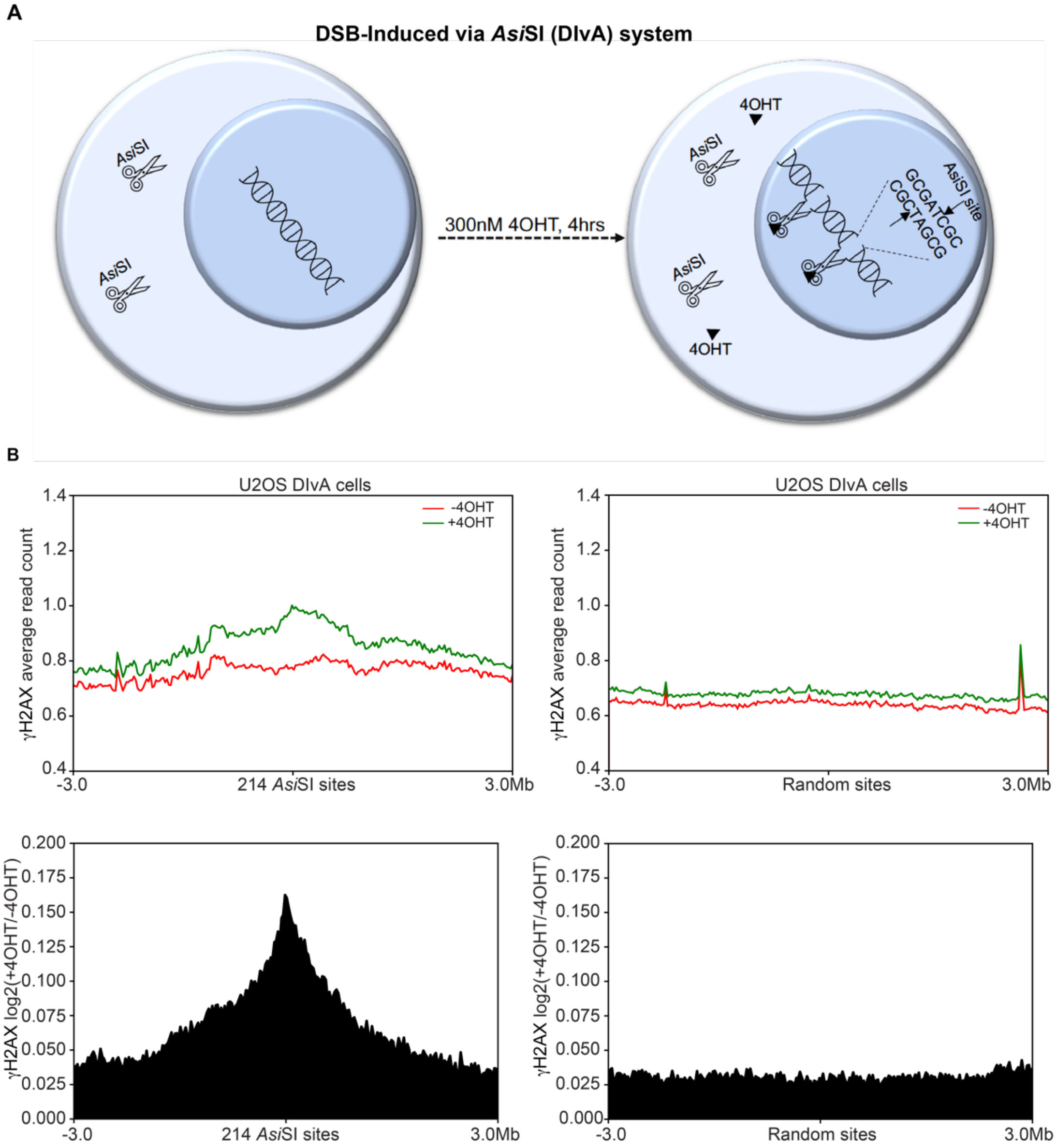


**Figure S1. Related to Figure 1: γH2AX enrichment at *AsiSI*-induced DSBs**
**(A)** A schematic describing DSB induction at *Asi*SI sites following 4OHT treatment in U2OS-DIvA cells. **(B)** Average profile of γH2AX enrichment between 4OHT-treated and untreated DIvA cells in 6Mb window surrounding 214 *Asi*SI sites with highest γH2AX (left) or 214 random genomic loci (right). Top: values are expressed as average normalized ChIP-seq reads. Bottom: values are expressed as average log2 ratio between 4OHT-treated and untreated cells.

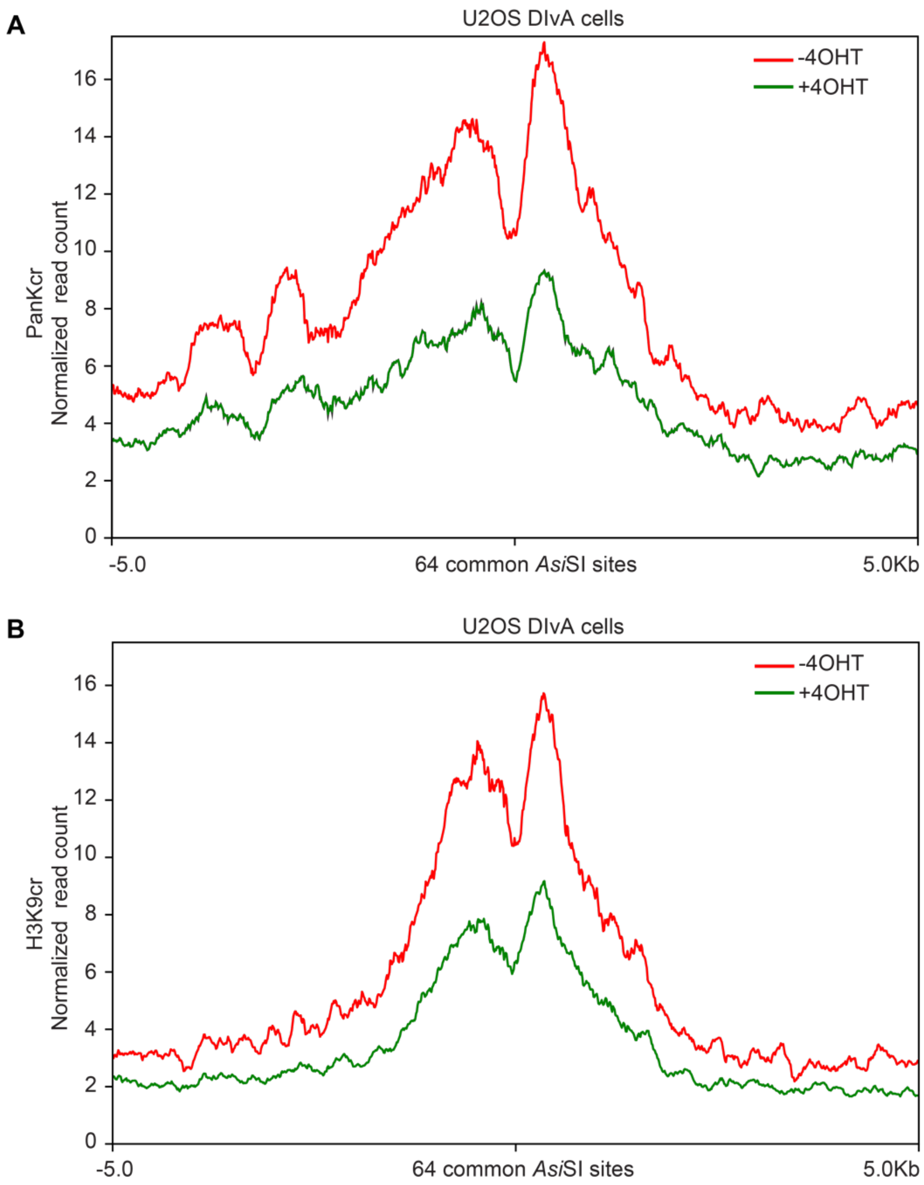


**Figure S2. Related to Figure 1: Reduction in PanKcr and H3K9cr at *Asi*SI-induced DSBs**
**(A-B)** Average normalized ChIP-seq reads of PanKcr (A) or H3K9cr (B) between 4OHT treated and untreated DIvA cells in 10kb window surrounding the 64 common cleaved *Asi*SI sites.

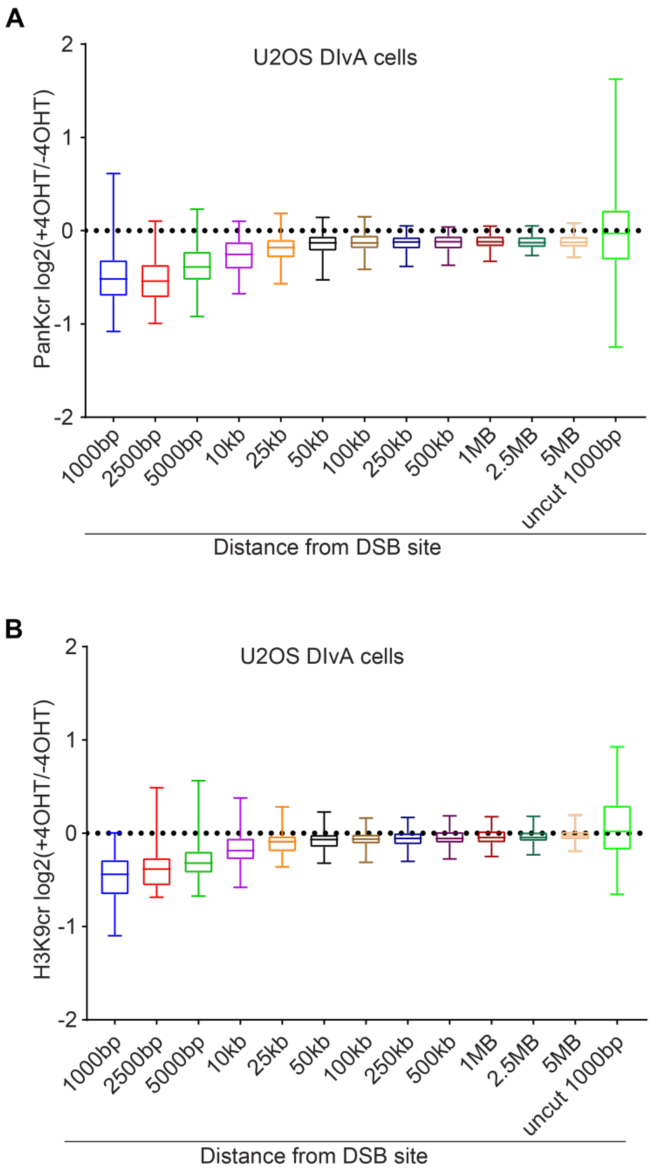


**Figure S3. Related to Figure 1: Distribution of PanKcr and H3K9cr surrounding *Asi*SI-induced DSBs**

**(A-B)** Boxplots representing the log2 ratio of PanKcr ChIP-seq (A) or H3K9cr ChIP-seq (B) between 4OHT-treated and untreated DIvA cells at the indicated distance surrounding the 64 common cleaved *Asi*SI sites.

A

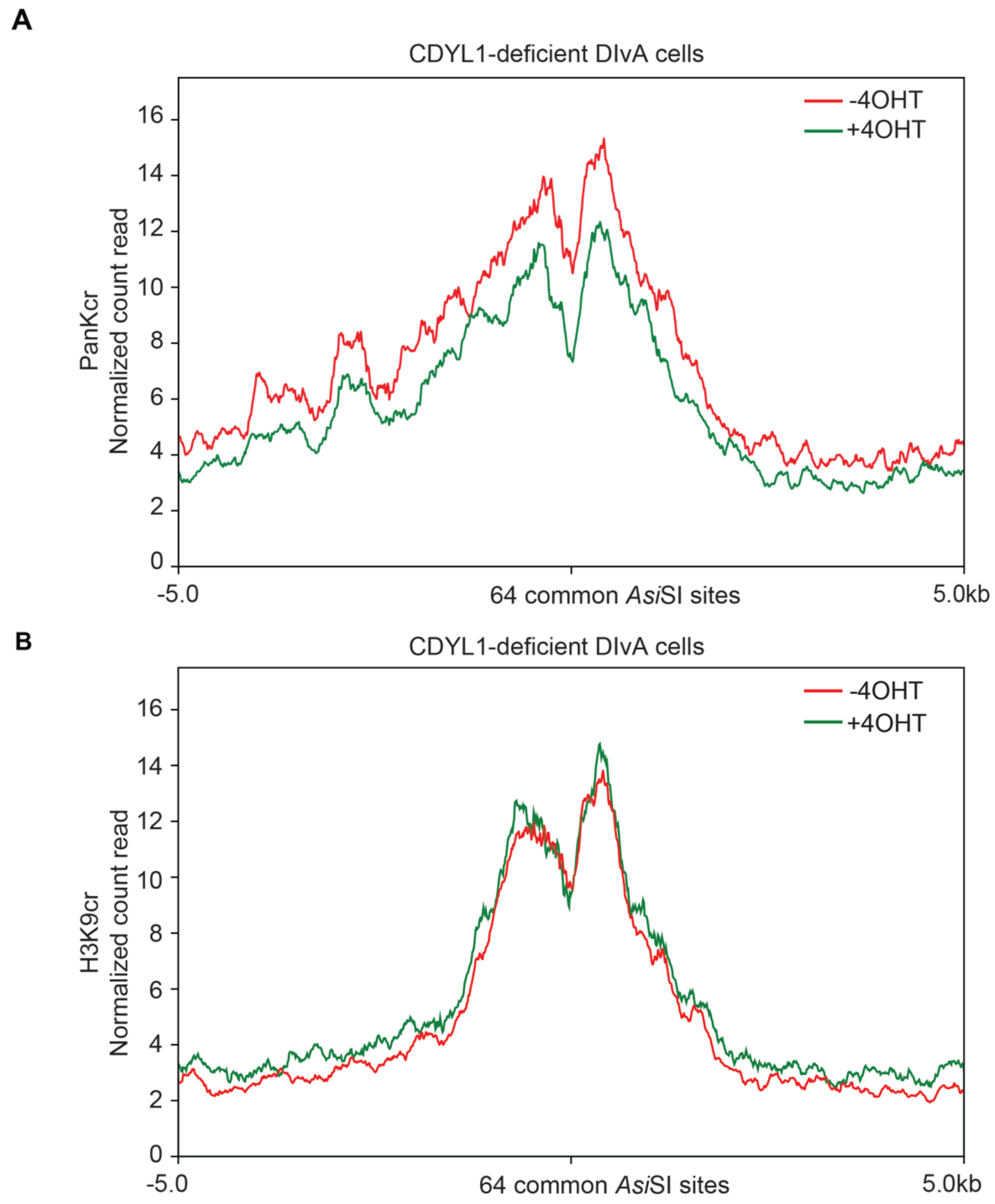


**Figure S4. Related to Figure 3: CDYL1-dependent reduction in PanKcr and H3K9cr at *Asi*SI-induced DSBs**

**(A-B)** Average normalized ChIP-seq reads of PanKcr (A) or H3K9cr (B) between 4OHT treated and untreated DIvA cells depleted of CDYL1 in 10kb window surrounding the 64 common cleaved *Asi*SI sites.

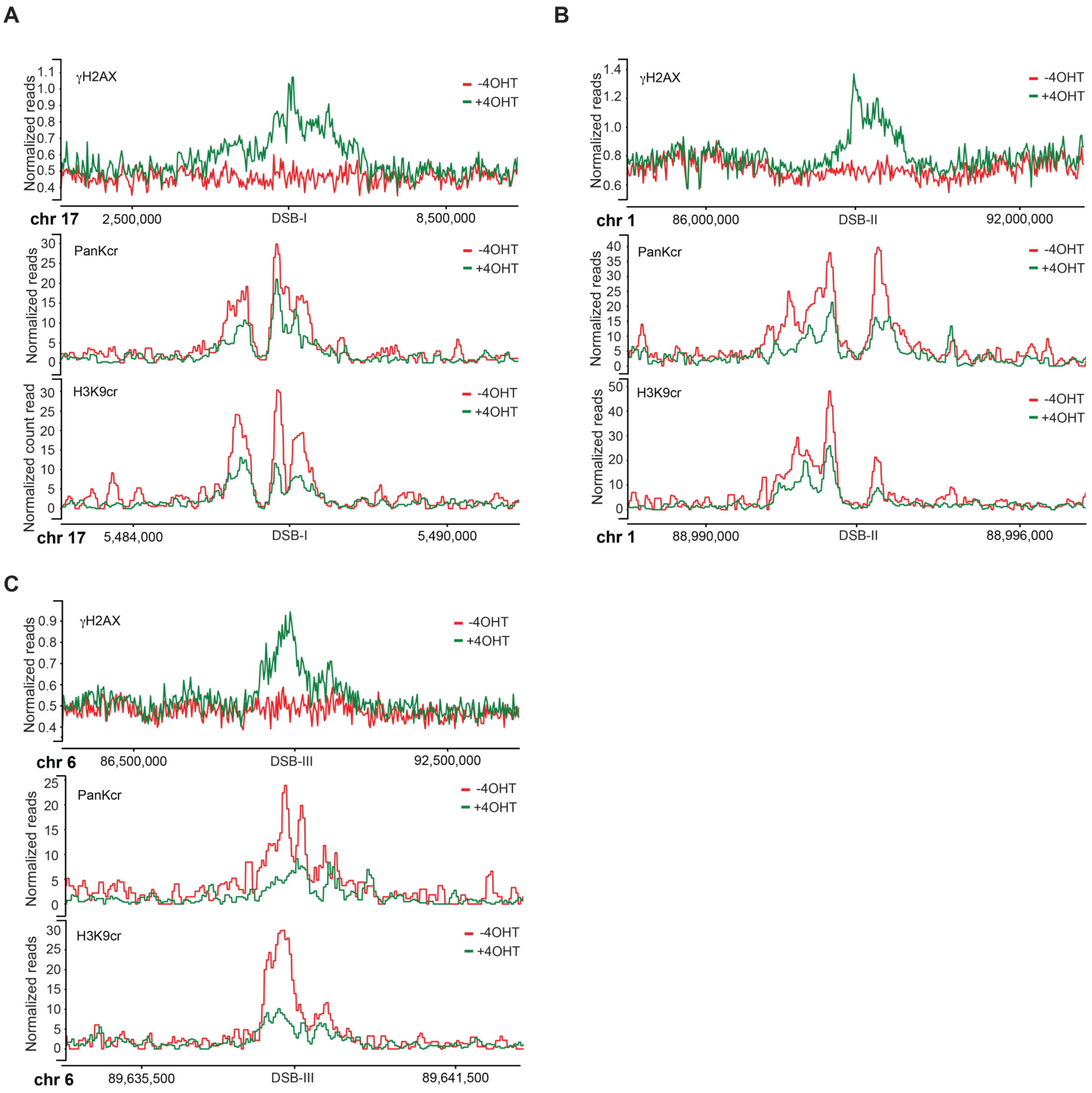


**Figure S5. Related to Figure 4: ChIP-seq profiles for 3 *Asi*SI-DSB sites**

**(A-C)** Genome Browser images representing average normalized ChIP-seq reads for γH2AX, PanKcr, and H3K9cr for three cleaved *Asi*SI sites: DSB-I (Chr17: 5,486,899-5,486,907) (A), DSB-II (Chr1: 88,992,912-88,992,920) (B), DSB-III (Chr6: 89,638,466-89,638,474) (C) in 4OHT-treated and untreated DIvA cells.

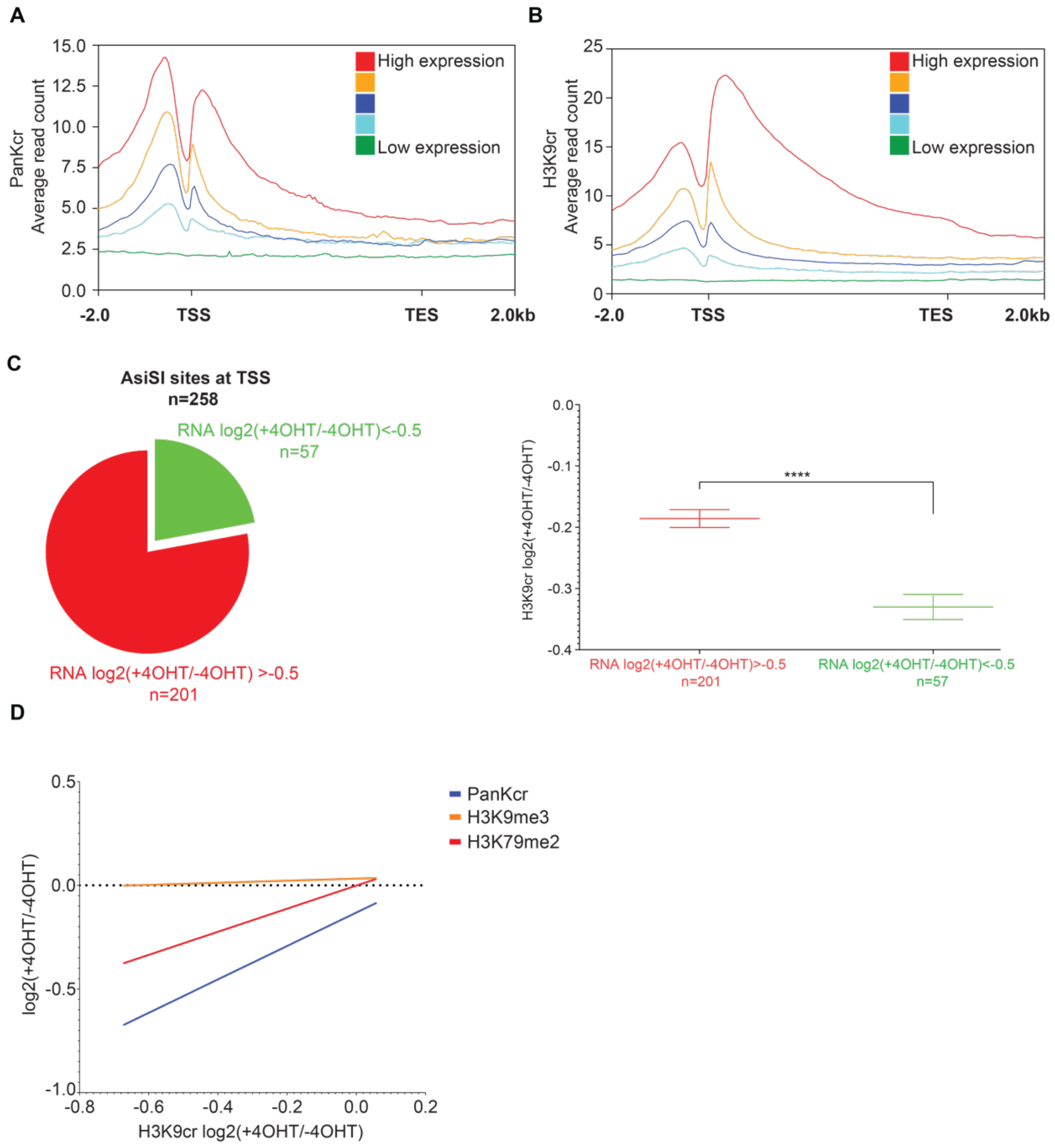


**Figure S6: Related to Figure 4: Reduction in H3K9cr correlates with DSB-induced transcriptional silencing**

**(A-B)** Average normalized ChIP-seq reads of PanKcr (A) and (H3K9cr) in untreated samples over human genes categorized by expression level based on previously published RNA-seq data of U2OS DIvA cells (Iannelli et al., 2017). **(C)** Left: Pie chart of 258 *Asi*SI sites at TSS classified by the reduction in gene transcription following 4OHT treatment based on previously published RNA-seq data (Iannelli et al., 2017). Red indicates genes with RNA log2(+4OHT/-4OHT)>-0.5 (n=201); Green indicates genes with RNA log2(+4OHT/-4OHT)<-0.5 (n=57). Right: Graph showing the mean log2 ratio of H3K9cr and standard error of the mean (SEM) between 4OHT-treated and untreated DIvA cells at 10kb window surrounding either 201 *Asi*SI sites (red) or 57 *Asi*SI sites (green) with the largest reduction in transcription. p-value was calculated using nonparametric Mann-Whitney test. *** p<0.0001. **(D)** Linear regression describing the relationship between H3K9cr profile (x-axis) and other histone modifications from previously published data (Clouaire et al., 2018) or PanKcr from this study (y-axis) around the 64 common cleaved *Asi*SI sites. Values are expressed as log2 ratio between 4OHT-treated and untreated DIvA cells.

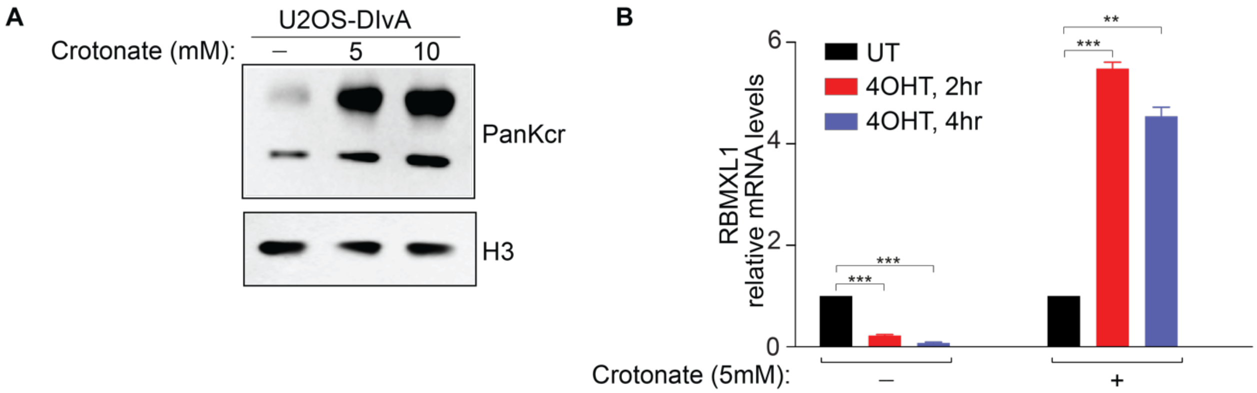


**Figure S7: Related to Figure 4: Elevated crotonyl-CoA levels impair DSB-induced transcriptional silencing**

**(A)** Western blot shows increased levels of PanKcr following the addition of the indicated concentrations of sodium crotonate. H3 is used as a loading control. **(B)** mRNA expression levels of RBMXL1 were measured before or after 300nM 4OHT treatment for 4hrs in DIvA cells that were either left untreated or pre-treated with 5mM sodium crotonate for 12hrs prior 4OHT treatment.

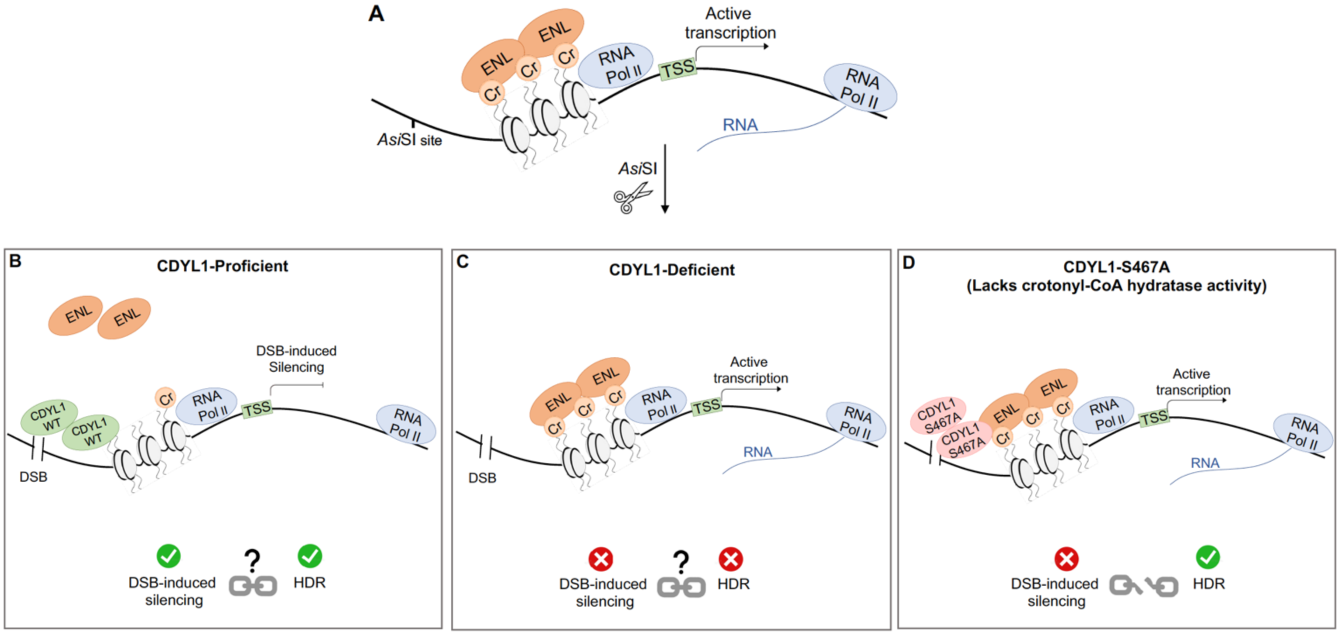


**Figure S8:** A hypothetical model showing that DSB-induced transcriptional silencing and HDR are functionally uncoupled. **(A)** Depicts an actively-transcribed gene nearby AsiSI recognition site in the absence of DSB. **(B)** CDYL1 is recruited to *Asi*SI-induced DSB to counteract Kcr, which fosters ENL eviction form TSS sites leading to DSB-induced transcriptional silencing and intact HDR. **(C)** CDYL1 depletion prevents Kcr reduction and ENL displacement from DSB sites and consequently leads to the alleviation of DSB-induced transcriptional silencing and defective HDR repair. **(D)** CDYL1 mutant that lost its crotonyl-CoA hydratase activity is recruited to DSB sites and foster HDR, but it is unable to counteract Kcr and elicit DSB-induced transcriptional silencing.

### Supplementary Tables

**Table 1: Reagent or Resource**

| | **Source** | **Identifier** |
|---|---|---|
| **Antibodies** | | |
| Anti-phospho-Histone H2A.X (Ser139), clone JBW301 | Millipore | 05-636 |
| Anti-β-actin [AC-15] (WB 1:10,000) | Sigma-Aldrich | A5441 |
| Anti-Histone H3 (WB 1:30,000) | Abcam | ab1791 |
| Anti-CDYL antibody (WB 1:1000) | Abcam | Ab5188 |
| Crotonyl-Histone H3 (Lys9) rabbit antibody (WB 1:1000) | PTM-BIO | PTM-516 |
| Anti-crotonyl lysine rabbit antibody (WB 1:1000) | PTM-BIO | PTM-501 |
| Anti ENL/MLLT1 | Bethyl | A302-267A |
| **Bacterial and Virus Strains** | | |
| DH5α Competent Cells | Thermo Scientific™ | EC0112 |
| ElectroMax Stbl4 | Invitrogen™ | 11635018 |
| **Chemicals and Recombinant Proteins** | | |
| Caffeine | Sigma-Aldrich | C8961 |
| ATMi | Sigma-Aldrich | KU-60019 |
| Ethidium bromide (EtBr) | Hylabs | BP451 |
| Polyethylenimine (PEI) | Polysciences | 343-6484 |
| Hexadimethrine bromide (Polybrene) | Sigma-Aldrich | H9268 |
| Puromycin | Invivogen | ant-pr |
| (z)-4-Hydroxytamoxifen | Sigma-Aldrich | H7904 |
| Crotonic acid | Sigma-Aldrich | 113018 |
| **Critical Commercial Assays** | | |
| WesternBright Quantom (ECL) | Advansta | K-12042 |
| TRIzol Reagent | Invitrogen | 15596026 |

| | | |
|---|---|---|
| qScript cDNA synthetsis kit | Quanta Bio | 95047 |
| Fast SYBR green master mix | Applied Biosystems | 4385610 |
| Phusion® High-Fidelity DNA Polymerase | NEB | M0530 |
| Protein A Magnetic beads | Invitrogen | 10002D |
| Protein G magnetic beads | Gene Script | L00274 |
| PFA | Electron Microscopy Science | 15710 |
| DNaseI kit | Promega | M6101 |
| MACHEREY-NEGEL Nucleospin kit | Ornat | 740609 |

**Software and Algorithms**

| | | |
|---|---|---|
| GraphPad Prism (v8.0) | GraphPad software lnc. | https://www.graphpad.com |
| ImageJ software (v1.8.0) | National Institutes of Health | https://imagej.nih.gov/ij/ |

**Table 2: Plasmids**

**Commercial plasmids**

| | | |
|---|---|---|
| pEGFP-C1 | Clontech Laboratories | #6084 |
| pDsRED-monomer-C1 | Clontech Laboratories | 632466 |
| pDsRED-monomer-CDYL1-WT | (Abu-Zhayia et al., 2018) | |
| pLKO.1 - TRC cloning vector | Addgene | #10878 |
| pMD2.G | Addgene | #12259 |
| psPAX2 | Addgene | #12260 |
| pX330-LMNA-gRNA1 | (Pinder et al., 2015) | N/A |
| pCR2.1-CloverLamin | (Pinder et al., 2015) | N/A |
| pX330-U6-Chimeric_BB-CBh-hSpCas9 | Addgene | #42230 |

**Plasmids generated in this study**

| Plasmid | Vector backbone | Insert |
|---|---|---|
| pEGFP-C1-CDYL1-S467A | pEGFP-C1-CDYL1-WT | All round PCR using primers F1,R1 |
| PLKO.1-TRC-Scramble | pLKO.1 - TRC cloning vector digested with EcoRI, AgeI | Annealed primers F2,R2 |
| PLKO.1-TRC-CDYL-shRNA | pLKO.1 - TRC cloning vector digested with EcoRI, AgeI | Annealed primers F3,R3 |

| | | |
|---|---|---|
| pX330-MIS12-gRNA1 | pX330-U6-Chimeric_BB-CBh-hSpCas9 cut with BbsI | Annealed Primers F4,F4 |
| pCR2.1 Clover-MIS12-LHA | pCR2.1 Clover-LMNA cut with BamHI, AfeI | PCR on genomic DNA extracted from U2OS-DiVA cells using primers F5,R5 and digested with BamHI, AfeI. |
| pCR2.1 Clover-MIS12 Donor | pCR2.1 Clover-MIS12-LHA cut with BsrGI, NotI. | PCR on genomic DNA extracted from U2OS-DiVA cells using primers F6,R6 and digested with BsrGI, NotI. |
| pDsRED-monomer-CDYL1-S467A | pDsRED-monomer-CDYL1 | All round PCR using primers F1,R1 |
| pLenti-IRES-CDYL-WT | pLenti-IRES-GFP Cut AgeI, PstI | PCR product from pEGFP-CDYL1 using primers F7,R7 |
| pLenti-IRES-CDYL-S467A | pLenti-IRES-GFP-CDYL-WT Cut PstI | Insert from pEGFP-C1-CDYL1-S467A digested with PstI |

**Table 3: Primers**

**Cloning Primers**

| Primer | Sequence |
|---|---|
| F1 | TGGCAAGGGCCTGGTTGCGCAGGTGTTTTGGCCCGGGACG |
| R1 | CGTCCCGGGCCAAAACACCTGCGCAACCAGGCCCTTGCCA |
| F2 | CCGGGTGGACTCTTGAAAGTACTATCTCGAGATTTGACGGGTGGATAATCTGTTTTT |
| R2 | AATTAAAAACAGATTATCCACCCGTCAAATCTCGAGATAGTACTTTCAAGAGTCCAC |
| F3 | CCGGGCTTAGGATTCATGCTGAGATCTCGAGATCTCAGCATGAATCCTAAGCTTTTTG |
| R3 | AATTCAAAAAGCTTAGGATTCATGCTGAGATCTCGAGATCTCAGCATGAATCCTAAGC |
| F4 | CACCGTGTGGATCCAATGACCTACG |
| R4 | AAACCGTAGGTCATTGGATCCACAC |
| F5 | GTAGGATCCCAGATGGCTTGGCTGGAGGACAAGCAAATTG |
| R5 | GGCTGAGCGCTCTTGGCTTATTTGCTATGTTGTTTTCAGTC |
| F6 | GCTGTACAAGAACTCTGTGGATCCAATGACCTACGAGGCC |
| R6 | AATGCGGCCGCAACAGTCACTGGTATGATTTGAACAGTCC |
| F7 | TAACCGGTATGGCTTCCGAGGAGCTGTACGAGGTTGA |

| | |
|---|---|
| R7 | ATCCTGCAGGTCAGAACTCATCGATCTTCCTCTGCAAG |

**ChIP Primers**

| Primer | Sequence |
|---|---|
| F1 (DSB-I) | GCAGAGAAGATGAGGCGGTAGAAG |
| R1 | CCAAGTAGCTGGGATTACAGGTGG |
| F2 (DSB-II) | ACTCAGGGAACTCCATTGGC |
| R2 | TACATCCGATTCGAGCCCTG |
| F3 (DSB-III) | AGGAATTGACTGCGGTGTTC |
| R3 | GGGGAGGAGGAAAGGTGTAG |

**Primers for Real time PCR**

| Primer | Sequence |
|---|---|
| MIS12-mRNA-F | GAGAGAAGATGAGGCGGTAGA |
| MIS12-mRNA-R | GCCAATGTCCTCAATTTGCT |
| RBMXL1-mRNA-F | TCAGGACTAGTTCGCAGCAG |
| RBMXL1-mRNA-R | TCGAGGTGGACCTCCATAAC |
| GAPDH-RT-F | CCAGGGCTGCTTTTAACTCT |
| GAPDH-RT-R | GGTGCCATGGAATTTGCCAT |
